# Even More Space Efficient Genome Sketching with Lash

**DOI:** 10.64898/2026.11.12.764074

**Authors:** Jianshu Zhao, Claire Wang, Yuhan Weng, Ruihang Xia, Antonio Gonzalez, Rob Knight

**Author notes:** Corresponding author: Jianshu Zhao or Rob Knight. Co-first author (Jianshu Zhao, Claire Wang).

## Abstract

**Motivation:** Advances in sequencing technology have driven an exponential growth in microbial genomic data, making exact pairwise distance computation increasingly impractical at scale. As a result, computationally efficient genomic distance estimation algorithms have become an essential alternative, e.g., sketching algorithms such as HyperLogLog and MinHash.

**Results:** Here, we present Lash, a software that provides UltraLogLog and ExaLogLog (more space-efficient variants of the HyperLogLog structure) and HyperMinHash (a more space-efficient variant of a MinHash structure) as methods to approximate genomic distance. UltraLogLog and ExaLogLog were demonstrated to be around 28% and 43% more space/memory-efficient respectively than HyperLogLog with the same accuracy, and HyperMinHash was 4-5 times more space/memory-efficient than MinHash for various genome comparison tasks.

**Availability and implementation:** Lash is released under the MIT license at https://github.com/jianshu93/lash.

## Introduction

There has been exponential growth in the amount of microbial genomic data with the improvement in high throughput sequencing technologies (Soon *et al*., 2013). Traditional large-scale genome analyses using exact distance computations are now extremely computationally expensive in time and memory due to the large amount of data. However, calculating the average nucleotide identity (ANI) (Ciufo *et al*., 2018; Richter *et al*., 2016; Jain *et al*., 2018; Richter and Rosselló-Móra, 2009), the distance between two genomes, is an important step in tasks such as classifying species, searching genomic databases, and clustering genomes. To deal with problems in scalability, alignment-free methods are becoming more popular. These are less accurate than traditional alignment methods such as BLAST but can scale to a larger number of genomes. Homologous-aware calculation is one type of alignment-free method based on finding homologous sequences via mapping-like tools, e.g., FastANI (Jain *et al*., 2018) and skani (Shaw and Yu, 2023). The other type is homology-unaware and relies on overall genomic similarity by using the Jaccard index 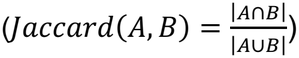 computed on sets of unique k-mers via sketching algorithms. This is then used to calculate ANI: 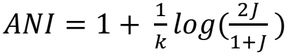. Probabilistic data structures (a.k.a sketching algorithms) are suited for this task and are used in many bioinformatics softwares: MinHash (Broder, 2002) via the software Mash (Ondov *et al*., 2016), and HyperLogLog (Flajolet *et al*., 2007) via the software Dashing (Baker and Langmead, 2019).

MinHash was applied for genomic distance estimation in Mash (Ondov *et al*., 2016) in 2016. In the original MinHash algorithm, each element is put into *p* independent hashing algorithms and the minimum hash values from each hashing algorithm are kept track of. The Jaccard similarity between two sets is then estimated as the fraction of the *m* positions at which the two signatures’ minimum values match. More optimizations have been made to MinHash, such as a one-permutation k-partition scheme (Li, A. Owen, *et al*., 2012), b bit minwise hashing (Li and König, 2010), densification (Shrivastava and Li, 2014) and optimal densification (Shrivastava, 2017). Bindash (Zhao, 2019) is a software for genome sketching and distance approximation that uses all the improvements from (Li and König, 2010); (Li, A. Owen, *et al*., 2012); (Shrivastava, 2017), together with the original MinHash (Broder, 2002), as well as a rolling hash. We chose Bindash as a reference to compare HyperMinHash (HMH) on genome sketching since it is the current SOTA MinHash algorithm. We also used its exact command as the ground truth for calculating exact Mash distances (1-ANI) to evaluate the accuracy of our other data structures.

HyperLogLog (Flajolet *et al*., 2007) (HLL) is a popular cardinality estimation method and has been applied to metagenome analysis previously in Dashing (Baker and Langmead, 2019) and KrakenUniq (Breitwieser *et al*., 2018). Extended HyperLogLog (Ohayon, 2021) is one of the improvements to HyperLogLog. In the original HyperLogLog (Flajolet *et al*., 2007), items are hashed to 64 bit binary numbers and mapped uniformly to registers. Each register contains 6 bits that track the maximum number of trailing 0s of all elements that have been hashed to the register. The harmonic mean, n, is computed on those values and the cardinality is estimated to be 2^n^.

As the size of genomic databases increases, there is a demand for space-efficient sketching of genomes so that memory requirement is not a bottleneck for making analyses. Here, in Lash, we implemented HyperMinHash (Yu and Weber, 2022) as a space efficient alternative of MinHash, and UltraLogLog (Ertl, 2023) and ExaLogLog (Ertl, 2024) as space efficient alternatives of HLL. Throughout this paper, we use space and memory interchangeably, since the size of a sketch determines both its storage on disk and its memory consumption while it is being built. We also refer directly to the sketch size as the memory requirement.

## Evaluation and Results

To show the space saving properties of UltraLogLog and ExaLogLog in our implementation (details can be found in the supplemental methods and materials section), we simulated sets with known cardinalities from 1 million to 4 million and evaluated space requirements (the size of sketches) for each algorithm and their corresponding precision parameter *P* against HyperLogLog. We found that when ULL (FGRA estimator) and ELL (ML estimator) have the same estimation error with HLL in terms of root mean square error (RMSE), ULL and ELL are 26.7% and 44.3% more space efficient than HLL, close to the theoretical 24.0% and 43.0% space savings (Fig. 1a). The results are consistent across several HLL precision *P* for the fixed 2 million cardinality (Fig.1b). We also implemented the HyperMinHash algorithm according to the original implementation in Python (details can be found in the supplemental methods and materials section). We also analyzed ULL’s improvements in memory variance product (MVP) from HLL and obtained a 24.3% and 28.4% improvement in ULL FGRA and ULL MLE, respectively (Supplementary Table S1). We calculated MVP by multiplying RMSE of the cardinality estimation (using simulated data), multiplied by the resulting sketch size. This aligns with findings in (Ertl, 2023).

**Figure 1.**
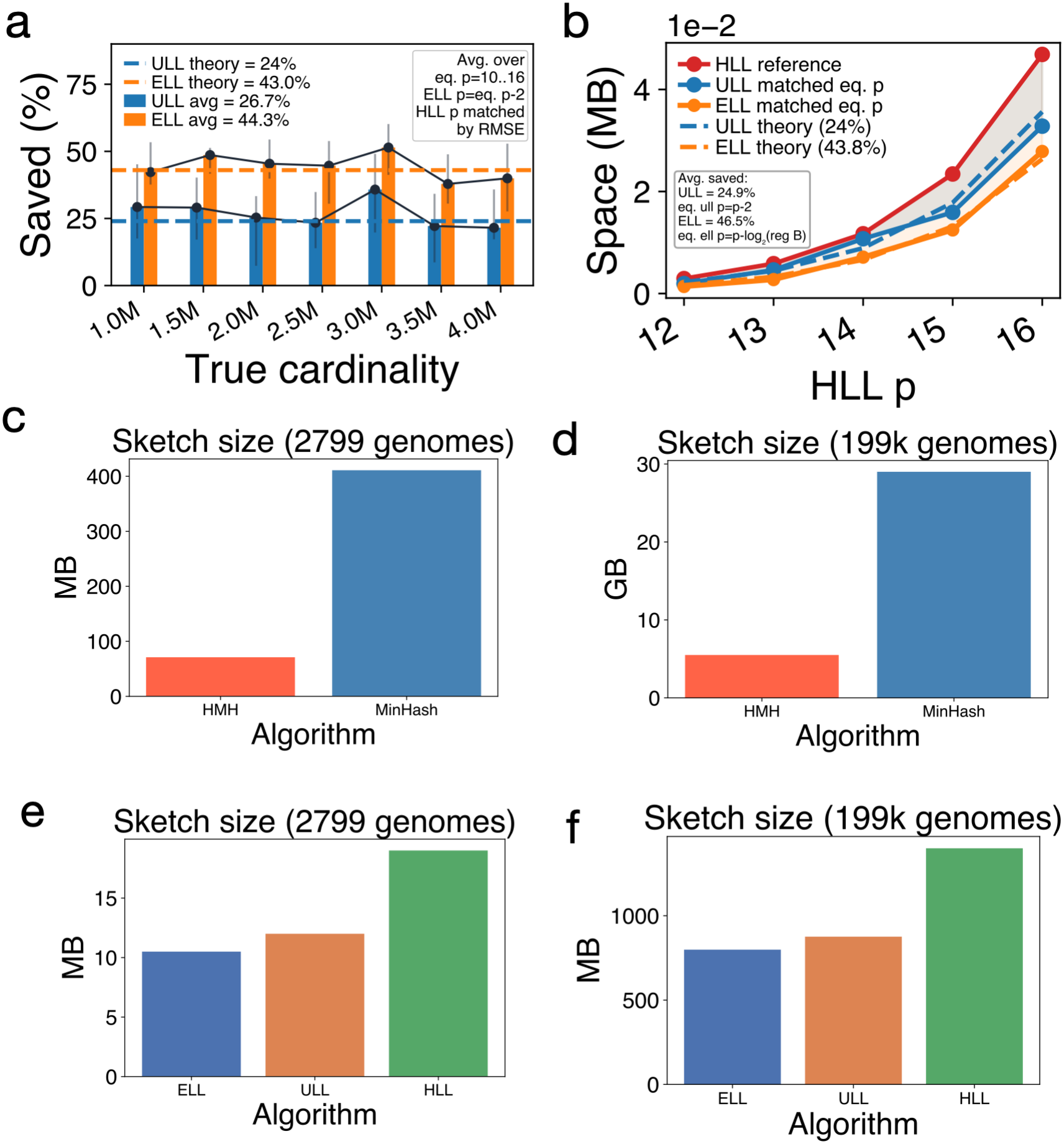
**a**. Space savings of ULL and ELL on simulated data across varied cardinalities. **b**. Space savings of ULL and ELL on simulated data with 2 million cardinalities with varied P. **c**. Sketch sizes of MinHash-based algorithms on 2799 genomes. **d**. Sketch sizes of MinHash-based algorithms on 199k genomes. **e**. Sketch sizes of HyperLogLog-based algorithms on 2799 genomes and **f**. Sketch sizes of HyperLogLog-based algorithms on 199k genomes.

We then compared the insertion time and estimation time for our implementation against the standard HyperLogLog. Consistent with the original paper, insertion for ULL is ∼3 times slower than HyperLogLog while ELL is 9∼10 times slower than HyperLogLog (Supplementary Fig. S2a). For estimation, ULL’s FGRA estimator is 3-4 times faster than the ULL ML estimator and around 2 times faster than the HLL default estimator (raw estimator) for genome cardinalities (1 million to 4 million distinct kmers) across *P* (Supplementary Fig. S2b). The ELL MLE estimator is the slowest, 20% to 60% slower than the ULL MLE estimator. For cardinality ranges from 10 to 10^7^, it has been shown that estimation time for ULL FGRA/ MLE and ELL MLE is very close with the same theoretical relative error (Ertl, 2023).

To make sure our findings transfer to genomic datasets, we benchmarked ULL, ELL, and HMH on 2 sets of genomic files: 2,799 *Streptomycetaceae* genomes and 199,923 representative GTDB genomes (Tan *et al*., 2026). We did not use Dashing 1.0 as a HLL baseline because we wanted a fair comparison using the same style implementation as a pure comparison of data structures using the same compiling language. Therefore, when we refer to HLL, we mean our own implementation of standard HLL within Lash following the same preprocessing instructions as the other algorithms. All our comparisons were run on 64 threads and k=16 (kmer length). As expected from theoretical analyses, HMH was demonstrated to be 4-5 times more space efficient than BinDash (Fig 1c-d), while ULL and ELL were demonstrated to be more space efficient than HLL by around 28% and 43% respectively (Fig 1e-f).

We also analyzed sketching and distance time on these genomic datasets. On 2799 *Streptomycetaceae* genomes, ELL, ULL, and HMH took 80.533, 73.726, and 88.811 seconds to run respectively, and HLL and MinHash (Bindash) took 75.769 and 31.599 seconds to run (Supplementary Fig S1a and S1c). On 199,923 GTDB genomes, ELL, ULL, and HMH took 2170, 2173, and 2160 minutes to run respectively, and HLL and MinHash (Bindash) took 2188 and 400 minutes to run respectively (Supplementary Fig S1b and S1d).

## Discussion and Conclusion

Space efficient sketching algorithms are becoming more important in modern genomics and metagenomics as we accumulate more microbial genomes (Shaw and Yu, 2023; Jain *et al*., 2018; Ondov *et al*., 2016; Zhao, 2019; Zhao, Zhao, *et al*., 2024). Here we further improve current practices in genome sketching methods by applying state of the art sketching algorithms. We showed that we can save >40% space (memory and disk space) for genome comparison tasks without losing accuracy via the SOTA LogLog-like algorithms such as ExaLogLog and HyperMinHash. ExaLogLog combines a space-efficient LogLog-style register design with maximum-likelihood estimation that asymptotically attains the Cramér–Rao lower bound for its register model, achieving an optimized memory–variance product (MVP) of approximately 3.67— about 43% less space or memory than standard HyperLogLog at equal accuracy (Ertl, 2024). Additionally, the ExaLogLog paper identifies compression as a route to further space savings without changing estimation accuracy; we use standard Zstandard (zstd), while specialized entropy coding remains a potential improvement. More recent efforts, such as Arithmetic Variable LogLog (Bushnell, 2026), can be more space-efficient than ExaLogLog. However, our implementation showed that its sketch is sensitive to insertion order (Supplementary Figure 3a-d), and that its merge operation is also sensitive to merge order (Supplementary Figure 4a-d). This is because AVLL discards rare high-rank history in the sketch. This makes it unsuitable for our use case, because pairwise genome distance calculation must be symmetric.

For microbial genome cardinalities of 1–5 million at *p*= 14, our HyperMinHash implementation uses the 6-bit rank for cardinality estimation, matching the original Python implementation’s leading-order HLL accuracy while incorporating Ertl’s improved range corrections (Ertl, 2017). Each rank–signature pair is packed into 16 bits (6 rank bits and 10 signature bits) and the full pair is used for similarity estimation, compared with 24 bits across separate arrays in the original Python implementation with the same configuration. Although the signature bits are currently excluded from our cardinality estimator, they contain additional cardinality information because register updates select among signatures when ranks tie. Maximum-likelihood estimation using the full register could therefore improve cardinality accuracy and potentially reduce memory requirements at a fixed cardinality error, although likelihood optimization may increase estimation time. This possibility is explicitly suggested in the ExaLogLog paper (HMH is a special case of ELL, which is ELL(*t*, 0)). Classic MinHash reduces Jaccard sampling error as the sketch size increases and converges to 0 (Li, A. B. Owen, *et al*., 2012); (Broder *et al*., 2000). HyperMinHash additionally introduces compression collisions whose contribution depends on signature width, while its collision correction relies on cardinality estimates whose attainable variance is constrained by the Cramér-Rao lower bound. Increasing register count (and therefore sketch size) reduces statistical uncertainty, but the Jaccard estimation error does not converge to 0.

The data structures and implementations introduced in Lash can be applied to many current bioinformatics tools and platforms that do genome clustering, genome database search, and large scale pairwise comparisons. For example, GSearch (Zhao, Both, *et al*., 2024) and gtdb-tk (Chaumeil *et al*., 2022) are database querying tools that require a distance metric to be computed while building an index, and dRep (Olm *et al*., 2017) relies on MinHash (via Mash) as the pre-clustering step. Because these tools store a sketch for every genome, smaller sketches would directly lower their memory usage. As genomic databases continue to grow, sketching methods with even more space efficiency will be of high practical relevance.

## Data and code availability

Lash is available on github at: https://github.com/jianshu93/lash, as a Rust crate at: https://crates.io/crates/lash-rs, on bioconda at https://anaconda.org/channels/bioconda/packages/lash/overview, and released on zenodo: 10.5281/zenodo.17412099. Lash integrates other open source Rust crates that we developed, including Hyperminhash, UltraLogLog and ExaLogLog, available via the Lash Github page. Our implementation of Arithmetic Variable LogLog is here: https://github.com/jianshu93/avloglog.

## Acknowledgements

We want to thank Otmar Ertl for his suggestions on the implementation of UltraLogLog and ExaLogLog.

## Author contributions

J.Z. designed the algorithm and implemented the key libraries. J.Z. and C.W. and implemented the Lash code. R.X. and J.Z. implemented the UltraLogLog library. J.Z. and C.W. wrote the manuscript. Y. W. and A. G. helped with the benchmark. R.K. supervised the work.

## References cited

Baker, D.N. and Langmead, B. (2019) Dashing: fast and accurate genomic distances with HyperLogLog. Genome Biol., 20, 265.

Breitwieser, F.P. et al. (2018) KrakenUniq: confident and fast metagenomics classification using unique k-mer counts. Genome Biol., 19, 198.

Broder, A.Z. et al. (2000) Min-wise independent permutations. J. Comput. Syst. Sci., 60, 630–659.

Broder, A.Z. (2002) On the resemblance and containment of documents. In, Proceedings. Compression and Complexity of SEQUENCES 1997 (Cat. No.97TB100171). IEEE Comput. Soc.

Bushnell, B. (2026) Arithmetic Variable LogLog: Advancing the memory-variance frontier. arXiv [cs.DS].

Chaumeil, P.-A. et al. (2022) GTDB-Tk v2: memory friendly classification with the genome taxonomy database. Bioinformatics, 38, 5315–5316.

Ciufo, S. et al. (2018) Using average nucleotide identity to improve taxonomic assignments in prokaryotic genomes at the NCBI. Int. J. Syst. Evol. Microbiol., 68, 2386–2392.

Ertl, O. (2024) ExaLogLog: Space-efficient and practical approximate distinct counting up to the Exa-scale. arXiv [cs.DS].

Ertl, O. (2017) New cardinality estimation algorithms for HyperLogLog sketches. arXiv [cs.DS].

Ertl, O. (2023) UltraLogLog: A practical and more space-efficient alternative to HyperLogLog for approximate distinct counting. arXiv [cs.DS].

Flajolet, P. et al. (2007) HyperLogLog: the analysis of a near-optimal cardinality estimation algorithm. Discrete Math. Theor. Comput. Sci., DMTCS Proceedings vol. AH,…

Jain, C. et al. (2018) High throughput ANI analysis of 90K prokaryotic genomes reveals clear species boundaries. Nat. Commun., 9, 5114.

Li, P., Owen, A.B., et al. (2012) One permutation hashing. In, Proceedings of the 26th International Conference on Neural Information Processing Systems - Volume 2, NIPS’12. Curran Associates Inc., Red Hook, NY, USA, pp. 3113–3121.

Li, P., Owen, A., et al. (2012) One permutation hashing for efficient search and learning. arXiv [cs.LG].

Li, P. and König, C. (2010) b-Bit minwise hashing. In, Proceedings of the 19th international conference on World wide web. ACM, New York, NY, USA.

Ohayon, T. (2021) ExtendedHyperLogLog: Analysis of a new Cardinality Estimator. arXiv [cs.DS].

Olm, M.R. et al. (2017) dRep: a tool for fast and accurate genomic comparisons that enables improved genome recovery from metagenomes through de-replication. ISME J., 11, 2864–2868.

Ondov, B.D. et al. (2016) Mash: fast genome and metagenome distance estimation using MinHash. Genome Biol., 17.

Richter, M. et al. (2016) JSpeciesWS: a web server for prokaryotic species circumscription based on pairwise genome comparison. Bioinformatics, 32, 929–931.

Richter, M. and Rosselló-Móra, R. (2009) Shifting the genomic gold standard for the prokaryotic species definition. Proc. Natl. Acad. Sci. U. S. A., 106, 19126–19131.

Shaw, J. and Yu, Y.W. (2023) Fast and robust metagenomic sequence comparison through sparse chaining with skani. Nat. Methods, 20, 1661–1665.

Shrivastava, A. (2017) Optimal densification for fast and accurate minwise hashing. ICML, abs/1703.04664, 3154–3163.

Shrivastava, A. and Li, P. (2014) Densifying one permutation hashing via rotation for fast near neighbor search. ICML, 557–565.

Soon, W.W. et al. (2013) High-throughput sequencing for biology and medicine. Mol. Syst. Biol., 9, 640.

Tan, B. et al. (2026) Phylogenomic survey of GTDB r232 expands the taxonomic and ecological range of alkylsuccinate synthase and distinguishes divergent AssA-like paralogs. Sci. Rep., 16.

Yu, Y.W. and Weber, G.M. (2022) HyperMinHash: MinHash in LogLog space. IEEE Trans Knowl Data Eng, 34, 328–339.

Zhao, J., Zhao, X., et al. (2024) BinDash 2.0: New MinHash scheme allows ultra-fast and accurate genome search and comparisons. bioRxiv.

Zhao, J., Both, J.P., et al. (2024) GSearch: ultra-fast and scalable genome search by combining K-mer hashing with hierarchical navigable small world graphs. Nucleic Acids Res., 52, e74.

Zhao, X. (2019) BinDash, software for fast genome distance estimation on a typical personal laptop. Bioinformatics, 35, 671–673.

