## Supplementary Material for "Even More Space Efficient Genome Sketching with Lash"

### Table of contents

Supplementary Table 1

Supplementary Figures 1-4

Supplementary Materials and Methods

### Supplementary Tables

**Supplementary Table S1**

| Estimator | Register memory | MVP | Reduction from HLL |
|---|---|---|---|
| HLL | 12 KB | 6.46 | - |
| ULL FGRA | 16 KB | 4.89 | 24.3% |
| ULL MLE | 16 KB | 4.63 | 28.4% |

MVP analysis for HLL and ULL (cardinality estimation on simulated data). Similar MVP analysis for ELL is in the original ELL paper.

### Supplementary Figures

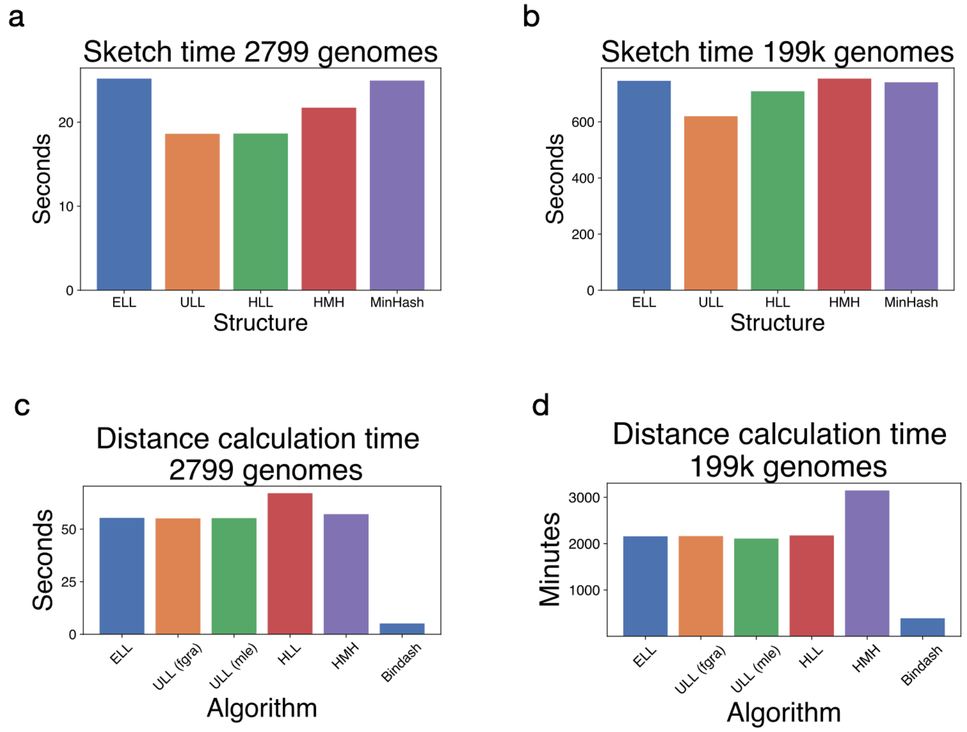


**Supplementary Figure 1**. **a.** Sketch time on 2799 genomes **b.** Sketch time on 199k genomes **c.** Distance calculation time on 2799 genomes **d.** Distance calculation time on 199k genomes

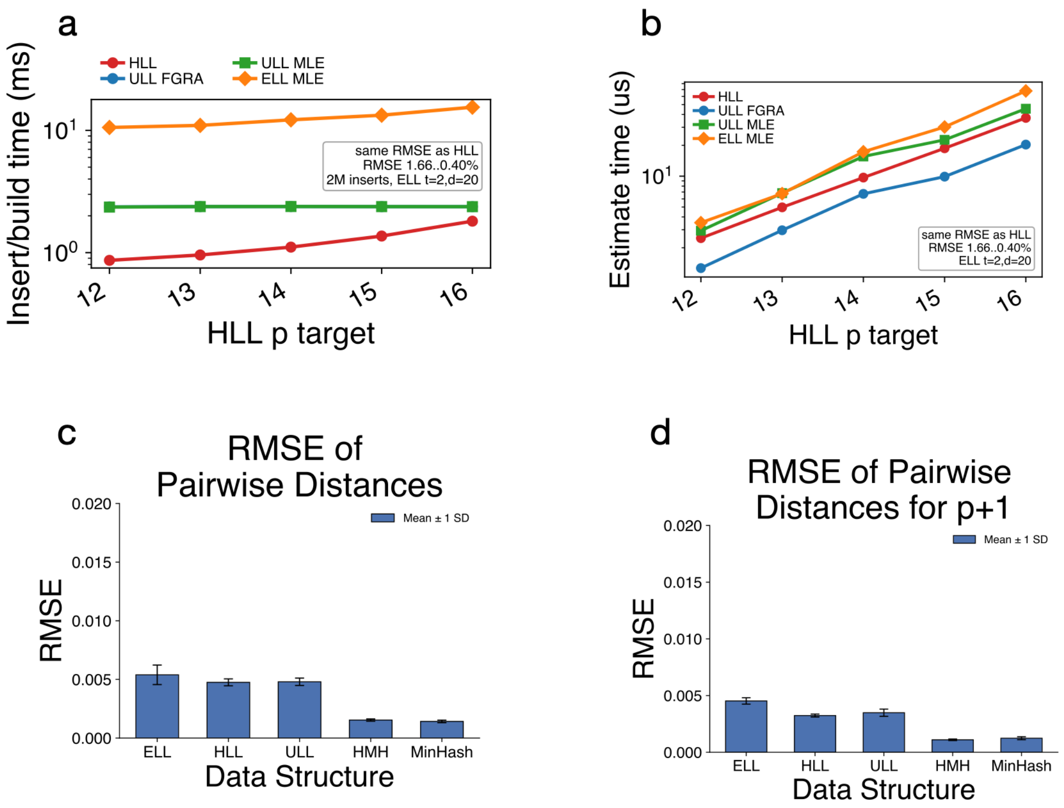


**Supplementary Figure 2. a.** Insertion/build time for LogLog algorithms (note ULL FGRA and ULL MLE have the same insertion procedure so they are both represented by the green line) **b.** Cardinality estimation time for LogLog algorithms **c.** RMSE of pairwise (1-ANI) distances for all algorithms **d.** RMSE of pairwise (1-ANI) distances for all algorithms using p + 1.

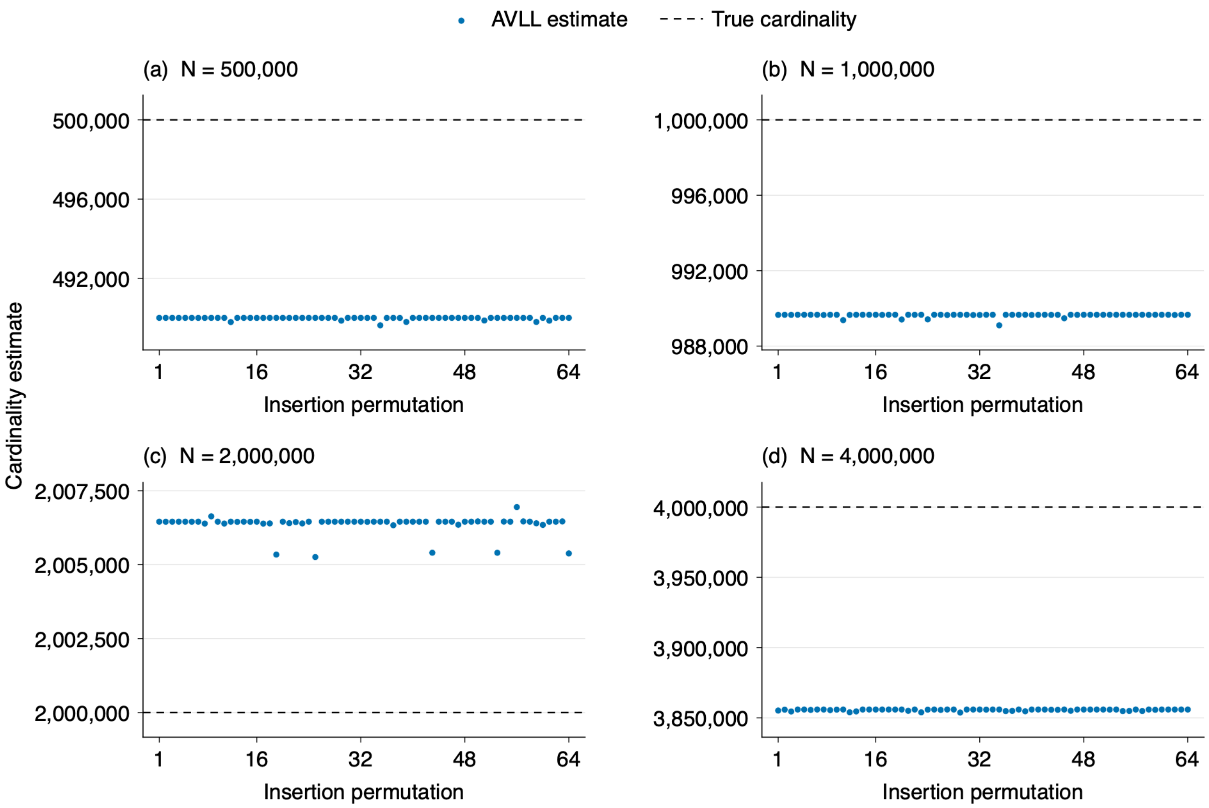


**Supplementary Figure 3.** AVLL estimates under 64 insertion permutations of fixed prefixes from one hash stream (seed 0). Panels (a)-(d) use true cardinalities N = 500,000, 1,000,000, 2,000,000, and 4,000,000, respectively. Every sketch contains 1,408 registers in a 1 KiB packed payload. Points show unmodified HLDLC estimates; dashed lines mark the exact cardinalities. The input set remains unchanged across permutations within each panel. These fixed-input results describe order sensitivity rather than population accuracy.

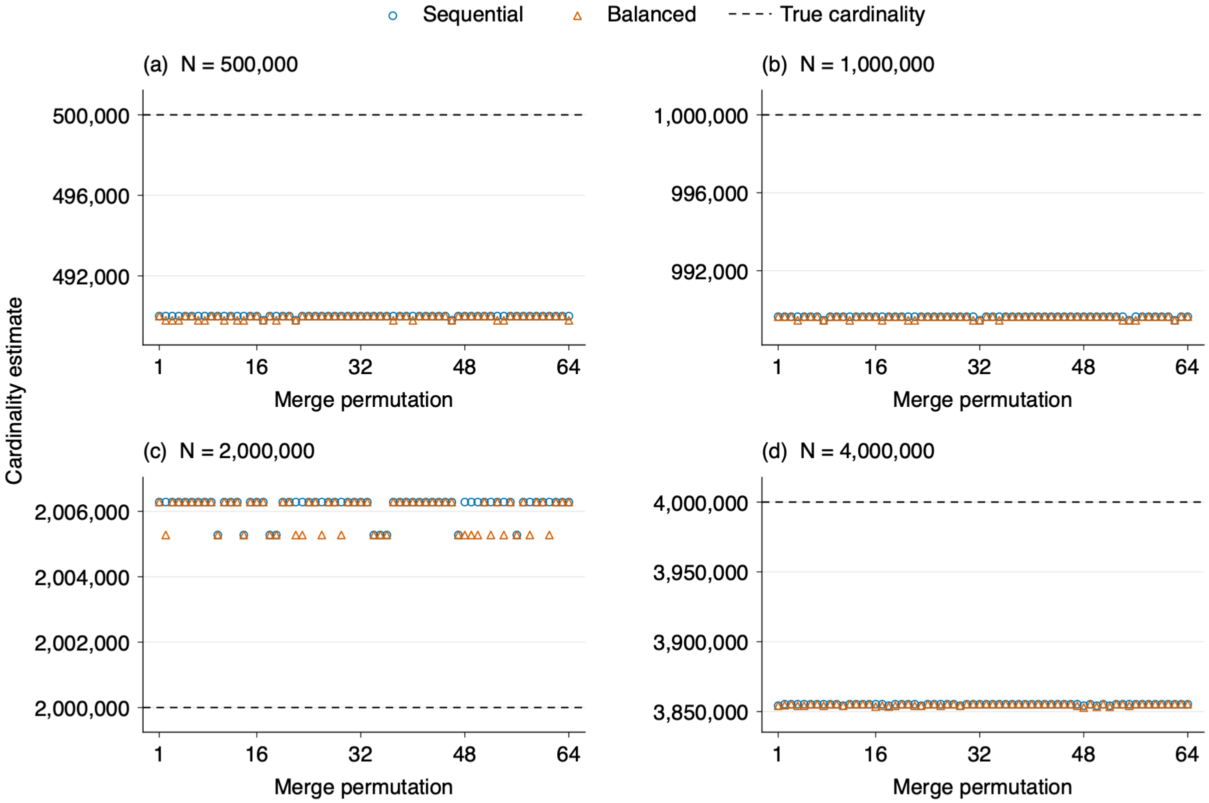


**Supplementary Figure 4.** AVLL estimates after merging eight fixed, disjoint shards under 64 shard permutations using sequential and balanced binary merge trees. Panels (a)-(d) use true cardinalities N = 500,000, 1,000,000, 2,000,000, and 4,000,000 from fixed prefixes of one hash stream (seed 0). Every sketch contains 1,408 registers in a 1 KiB packed payload. Points show unmodified HLDLC estimates after all shards have been merged; dashed lines mark the exact cardinalities. Shard membership and within-shard insertion order remain fixed. These results describe merge-order and tree effects for these inputs rather than population accuracy.

<u>Materials and Methods</u>

**HyperMinHash implementation**. Our Rust implementation of HyperMinHash (HMH) follows the space-efficient sketching framework of Yu and Weber (2022) to estimate set cardinality and Jaccard similarity. HMH uses one-permutation partitioning to distribute input elements, such as encoded k-mers. Among $m = 2^p$ buckets, HMH maintains one compact representative per bucket rather than the globally smallest $m$ hashes. By default, our implementation hashes each input element using XXH3-128, although externally computed 128-bit hashes are also supported. The upper $p$ bits of the hash's lower 64-bit word determine the bucket index. Each bucket contains a 16-bit register comprising a 6-bit rank and a 10-bit signature, with zero indicating an empty bucket. The rank records the number of leading zeros after the bucket-index bits plus one, so a suffix beginning with 001 has rank 3. Unlike the paper's literal floating-point construction, which retains mantissa bits immediately following the first nonzero bit, our implementation obtains its signature from the lowest 10 bits of the hash's upper 64-bit word. Each update retains the larger packed register value, prioritizing a larger rank and, when ranks are equal, a larger signature. This rank-signature representation preserves the HyperLogLog-inspired compression principle while using a different representative ordering from the paper's minimum-mantissa convention. The bucket precision is configurable from $p = 4$ to $p = 26$, with a default of $p = 14$, corresponding to 16,384 registers and 32 KiB of register storage. Jaccard estimation counts equal, nonzero registers and excludes buckets empty in both sketches from the denominator. For nonidentical sketches, the default estimator subtracts an estimated number of accidental register matches before normalization and bounds the result to $[0,1]$, while identical sketches receive similarity 1. This collision correction is approximate, and the optional fast estimator omits it; consequently, the uncorrected fraction of matching compressed registers should not be equated directly with the Jaccard index. Cardinalities are estimated from register ranks using Ertl's improved raw HyperLogLog estimator (Ertl, 2017), which is an additional implementation choice within the HMH framework. Compatible sketches are merged by taking register-wise maxima, and intersection cardinality is estimated by multiplying corrected Jaccard similarity by estimated union cardinality, capped at the smaller estimated input cardinality.

**UltraLogLog implementation.** Our Rust implementation of UltraLogLog (ULL) follows Ertl's Java implementation. Input elements are hashed using XXH3-64 by default, with support for alternative hash functions or precomputed 64-bit hashes. The sketch comprises $m = 2^p$ initially zero-valued, 8-bit registers, with configurable precision $3 \leq p \leq 26$ and a register-storage requirement of $m$ bytes. The most significant $p$ hash bits select a register, while the number of leading zeros in the remaining bits determines a geometrically distributed update rank. Each register combines a 6-bit encoding of the maximum observed rank with two additional bits that record whether ranks one and two below that maximum have also occurred. These additional bits preserve information discarded by HyperLogLog, improving the statistical efficiency of cardinality estimation. Following the paper's practical encoding, our implementation updates a register by unpacking its retained rank information into a bit mask, setting the bit corresponding to the incoming rank, and repacking the result into one byte. Updates take constant time, are independent of insertion order, and are unaffected by repeated insertion of the same element. Sketches constructed with consistent hashing can be merged by combining their retained rank information, with support for reducing precision and merging sketches at the lower of their two precisions. The default cardinality estimator is the optimal further generalized remaining area (FGRA) estimator, which aggregates contributions from the register states using precomputed constants and incorporates corrections for small and large cardinalities. An optional maximum likelihood estimator (MLE) numerically solves the likelihood equation and applies a finite-register bias correction. Although generalized remaining area (GRA) estimation (Wang and Pettie, 2023) provides the methodological foundation for FGRA, the implementation uses FGRA and MLE rather than a separate GRA estimator. Both estimators directly return an estimate $\hat{n}$ of the distinct count $n$, without an additional doubling or exponentiation step. The paper reports approximately 28% lower memory requirements than densely packed 6-bit HyperLogLog when using maximum likelihood estimation, or approximately 24% lower requirements with FGRA, at comparable asymptotic estimation accuracy. These savings describe the memory-accuracy trade-off and do not imply that using exactly half as many registers always provides the same accuracy.

**ExaLogLog implementation**. Our Rust implementation of ExaLogLog (ELL) follows Ertl 2025 for approximate distinct counting, including applications to sets of encoded k-mers. Input elements are hashed using XXH3-64 by default, with support for alternative hash functions and precomputed 64-bit hashes. The sketch comprises $m = 2^p$ initially zero-valued registers, each containing $6 + t + d$ bits, where $t$ controls the resolution of update values and $d$ determines how many smaller update values are tracked. Each hash is divided into its lowest $t$ bits, the next $p$ bits specifying the register index, and the remaining upper bits used to count leading zeros. The update value is calculated as $k = 2^t L + j + 1$, where $L$ is the leading-zero count and $j$ is the integer represented by the lowest $t$ bits. This construction provides a computationally inexpensive approximation to a geometric distribution with finer resolution than the update distribution used by HyperLogLog and UltraLogLog when $t > 0$. Each register stores its largest observed update value $u$ in $6 + t$ bits and uses a $d$-bit bitmap to indicate which values from $u - d$ through $u - 1$ have occurred. When a larger value arrives, the bitmap is shifted to align with the new maximum; smaller values within the retained range set their corresponding bits. These updates take constant time, are independent of insertion order, and do not change the sketch when an element is inserted repeatedly. Our implementation stores registers densely in a byte vector, supports configurable $t$, $d$, and $p$, and provides specialized access for 32-bit registers. Cardinality is estimated using the paper's maximum likelihood approach, with numerical solution of the likelihood equation followed by the parameter-dependent bias correction $\hat{n} = \hat{n}_{ML}/[1 + c(t, d)/m]$. Compatible sketches using consistent hashing and the same $t$ can be merged, with the resulting sketch retaining the smaller $p$ and $d$ of the inputs. The implementation also supports 32-bit hash tokens, but does not provide the paper's separate sparse-mode estimator or martingale estimator. The paper's approximately 43% memory reduction relative to densely packed 6-bit HyperLogLog applies to the optimized configuration $t = 2, d = 20$, which uses 28-bit registers, at comparable asymptotic estimation accuracy. This configuration is supported by our implementation, while other parameter choices produce different memory–accuracy trade-offs

<u>Sketching</u>

In the sketching process of Lash, each canonical k-mer is first compressed into 2 bits per nucleotide (A=00, C=01, G=10, T=11), with any unused bits masked by 0s. This varies per value of k; our iterator for sequences accepts 32 bits or 64 bits so any value of k besides 16 or 32 results in masking. The encoded k-mer is then hashed with XXH3-128 into a 128-bit integer for HyperMinHash or a 64-bit integer for the LogLog algorithms. A subset of $p$ hash bits selects one of $m = 2^p$ registers (called buckets in HyperMinHash), and the remaining bits determine the value used to update that register. Each algorithm keeps different information per register. In HyperMinHash, each bucket holds a 16-bit register made of a 6-bit rank (leading zeros plus one) and a 10-bit signature, and an update retains the larger packed value. In the LogLog family (HLL, ULL, ELL), the register keeps the maximum rank observed, based on the number of leading zeros. UltraLogLog adds two bits per register recording whether the two ranks below the maximum have also occurred, and ExaLogLog stores a d-bit bitmap of smaller update values, which HyperLogLog discards. The structure is then compressed with zstd and serialized to a sketch file on disk.

In all sketching algorithms, the precision p can be specified, which sets the number of registers ($2^p$). A higher p gives more accurate distance computation, since more registers make the estimate more robust to registers with outlier values, but it also uses more memory. For each data structure, the sketch size approximately doubles each time p increases by 1 (e.g., the default p = 14 for HyperMinHash gives 16,384 registers, or 32 KiB of register storage).

The sketch step was parallelized via Rayon across each input genome/file. Users input a file path, which contains paths to multiple genomes, and all genomes are constructed into one sketch.

<u>Distance</u>

For the LogLog sketches (ULL and ELL), distance is calculated by merging sketches and estimating the cardinalities of the union and intersection (see the Jaccard equation). To form the union, corresponding registers from the two sketches are paired and their retained information is combined, keeping the larger value in each register. Merging requires consistent hashing, and sketches with different precisions are merged at the lower one, although in practice we sketch reference and query sets with the same precision. Registers are treated as independent and identically distributed under a Poisson model of the number of distinct elements, so the estimators pool information across all registers to estimate the distinct count n directly. For ULL, this is done with the FGRA estimator (default) or an optional MLE. For ELL, it is done with MLE followed by a parameter-dependent bias correction. The intersection is not estimated directly but is obtained by inclusion-exclusion: $|A \cap B| = |A| + |B| - |A \cup B|$.

Using the three cardinalities ($|A|, |B|, |A \cup B|$), the Jaccard similarity $\frac{|A \cap B|}{|A \cup B|}$ is calculated, and from it the final metric, 1 − ANI. There are two possible equations for ANI according to different evolutionary models: $ANI = 1 + \frac{1}{k} ln(\frac{2J}{1+J})$ and $ANI = (\frac{2J}{1+J})^{\frac{1}{k}}$. By default, Lash uses the first one.

For HyperMinHash, similarity is instead computed directly from the registers. The default estimator counts equal, nonzero registers (excluding buckets empty in both sketches), subtracts an estimated number of accidental matches, and bounds the result to [0, 1]. Identical sketches receive J = 1, and an optional fast estimator skips the collision correction. Union cardinality comes from the register-wise maximum merge and Ertl's improved raw HyperLogLog estimator, and intersection is estimated as J times the union cardinality, capped at the smaller input cardinality. ANI and 1 − ANI are then computed as above.

The distance step was parallelized via Rayon across each reference file/genome. Users input a reference and query file prefix to the file names of their respective sketches, and all pairwise distances are calculated.

<u>Benchmark details</u>

In order to compare HMH's space savings from BinDash, we ran BinDash with different sketch sizes (controlled by the number of bins) and compared it against the RMSE of HMH on 60 sampled files. We found that BinDash with 19200 bins (sketch_size=300) had a similar RMSE to HMH on p=14, and we use this version to compare memory and speed for a fair comparison.

The set of 199,923 genomic files was selected from GTDB R232's 199,923 representative species per cluster (Tan *et al.*, 2026). The set of 2799 genomic files was selected from a collection of the bacterial family *Streptomycetaceae*. Accuracy benchmarks were run on a set of 60 genomic files selected from those 2799 genomic files and their exact pairwise distances. Similar RMSE was achieved across algorithms in the same category (LogLog or MinHash-like) (Supplementary Fig S2c). The RMSE is also shown for both our p of choice and p+1 (Supplementary Fig S2c-d).

Our implementation of ELL with *t*=2, *d*=20 is the theoretical maximum space saving achievable by ExaLogLog according to the Cramér-rao bound which states that the theoretical MVP (memory variance product) for an unbiased estimator is MVP $\approx \frac{(q+d)\ ln\ b}{\zeta(2,1+\frac{b^{-d}}{b-1})}$, where $\zeta$ is the Hurvitz zeta function.

All benchmarks were run on 64 threads on an AMD EPYC 7302 32-Core Processor.

<u>References</u>

Ertl,O. (2017) New Cardinality Estimation Methods for HyperLogLog Sketches. *arXiv [cs.DS]*.

Tan,B. *et al.* (2026) Phylogenomic survey of GTDB r232 expands the taxonomic and ecological range of alkylsuccinate synthase and distinguishes divergent AssA-like paralogs. *Sci. Rep.*, **16**.

Wang,D. and Pettie,S. (2023) Better Cardinality Estimators for HyperLogLog, PCSA, and

Beyond. In, *Proceedings of the 42nd ACM SIGMOD-SIGACT-SIGAI Symposium on Principles of Database Systems*. ACM, New York, NY, USA.
